# Unexpected softening of giant unilamellar vesicles by budding yeast septin filaments: a curvature dependent mechanism

**DOI:** 10.64898/2026.04.27.721050

**Authors:** Brieuc Chauvin, Luca Costa, Martin Lenz, Bassam Hajj, Pierre-Emmanuel Milhiet, Stéphanie Mangenot, Aurélie Bertin

## Abstract

Budding yeast septins assemble into filamentous networks bound to the inner plasma membrane. *In situ* or *in vitro*, septins are implicated in membrane deformations. We therefore suspected that septins might alter membrane mechanical properties both directly or indirectly. To decipher whether septins directly tune the rigidity of membranes, we used a cell free *in vitro* approach. To this end, using AFM, we measured the mechanical response of reconstituted GUVs pre-incubated with septins. Unexpectedly, we find that large GUVs (typically tens of µm diameter size) are more deformable in the presence of septins. Theoretical modeling suggests that this peculiar behavior is likely due to initial micrometer membrane “wrinkled” deformations imposed by septins. Conversely, small GUVs (1 to 2 microns in diameter) cannot undergo any micrometric deformations and are thereby less deformable with septin filaments bound. Our findings suggest that, in specific cellular context, septins could provide a membrane reservoir and eventually facilitate membrane deformations.

**Significance statement:** Filamentous cytoskeletal septins, interacting with membranes would be expected to enhance membrane rigidity. Upon mechanical stress, GUVs larger than tens of microns appear, more deformable in the presence of septins. Septins’ initial membrane reshaping is responsible for this unexpected behavior, as shown by theoretical modeling. However smaller non deformable vesicles are more rigid, with septins bound.

## Introduction

Cell membranes need to undergo constant reshaping with cells dividing, migrating, achieving endocytosis, exocytosis among other membrane remodeling events. Within cells, the rigidity of membranes varies both in time and space. The formation of filopodia or lamellipodia requires fast local modifications in membrane mechanics from being soft to rigid. Conversely, some cellular compartments or domains (cleavage furrow during cell division) must be rigid for a larger time scale. Membrane rigidity is known to be controlled by the activity of the actin cortex interacting with the membrane. The membrane itself can only be stretched by a few percent. However, the cortex can induce the formation of protrusions and invaginations which provide a membrane reservoir that can be remodeled under tension (Raucher and Sheetz, 1999). Well-characterized proteins (Bar proteins, spectrins) linking the membrane and the actin cortex were shown to be involved in controlling the rigidity of the plasma membrane (Arkhipov *et al*., 2008; Legendre *et al*., 2008). Septins are cytoskeletal proteins able to tune the mechanical properties and the shape of membranes (Gilden and Krummel, 2010). They are ubiquitous in eukaryotes and are protein complexes self-assembling into palindromic hexamers or octamers gathering two copies of three or four different subunits (Mendonça *et al*., 2019). Septins can further assemble into filamentous structures either in solution or bound to membranes (bundles, rings, arrays of filaments), both *in situ* and *in vitro* (Kinoshita, 2003; Garcia *et al*., 2011; Taveneau *et al*., 2020). They were shown to interact specifically with phosphoinositides (Zhang *et al*., 1999; Bertin *et al*., 2010; Beber *et al*., 2019a). They also interact directly with actin (Mavrakis *et al*., 2014) and could thus connect the membrane to the cortex. Septins are often co-localized with actomyosin contractile rings, specifically during cytokinesis (Spiliotis, 2018). In cellular contexts, they appear to stiffen membranes and stabilize the cortex. For instance, septins are localized at the annulus in spermatozoa. Septin depletion (Sept4) leads to the generation of a sharp bend or rupture at the annulus (Ihara *et al*., 2005). In motile T cells, septin depletion induce spontaneous blebbing and protrusions and thereby uncontrolled motility (Tooley *et al*., 2009). Within those cells, septins are colocalized with the actin cortex. Interestingly, septins were shown to facilitate membrane fusion and facilitate macropinosome maturation and subsequent fusion with endosomes and lysosomes (Dolat and Spiliotis, 2016). Septins are involved in endocytosis events (Krauß and Haucke, 2007) possibly through their interaction with dynamins (Maimaitiyiming *et al*., 2013), and phagocytic uptake (Huang *et al*., 2008). Septins have recently been pointed as mechano-protective and mechanosensitive cytoskeletal elements (Mayca-Pozo *et al*., 2026; Utgaard *et al*., 2026). Their mechanotransducive mechanism however remains to be elucidated.

In that context, we have quantitatively probed how septins can directly affect the mechanical properties of membranes. To decouple the role of septins from those of their partners actin and myosin, we have carried out bottom-up cell-free assays using a controlled set of components and parameters. Deformable Giant Unilamellar Vesicles (GUVs) were employed. In comparison with supported lipid bilayers, GUVs’ membranes are free standing and unaffected by the presence of substrates. Septins do deform GUVs in a periodic manner because they are curvature sensitive (Tanaka-Takiguchi *et al*., 2009; Beber *et al*., 2019b; Nakazawa *et al*., 2022). Unexpectedly, using GUVs and micropipettes manipulation, we were able to show that the bending modulus and elastic area modulus of biomimetic membranes were barely affected by the presence of septins (Beber *et al*., 2019b). Here, we assess the global mechanical properties of GUVs decorated with septins using AFM force spectroscopy. The Young’s modulus of supported lipid bilayers can be estimated using AFM (Butt and Franz, 2002a; Fonseka *et al*., 2022). AFM spectroscopy has also been widely used to estimate the mechanics of adherent cells (Gavara and Chadwick, 2012; Krieg *et al*., 2019), interaction forces between cells or biomimetic lipid bilayers and components of supramolecular assemblies, as well as intramolecular forces stabilizing the fold of a protein (Baumgartner *et al*., 2000).

Our investigations show that overall, GUVs reshaped by septins at micrometric scales appear more deformable than without septins bound. Septins generate a membrane reservoir which is stretched when a force is applied with an AFM probe. Conversely, smaller vesicles (below 3 µm in diameter) appear stiffer with septin bound since no prior deformations can be generated.

## Results

### Septins restrict micropipette aspiration of GUVs

To decipher whether septins affect the mechanical properties of membranes, we set up micropipette aspiration experiments (Shi and Baumgart, 2015). Briefly, a micropipette connected to a pressure controller is brought in close vicinity of a GUVs. Upon aspiration, the vesicle is deformed and a tongue is generated inside the micropipette. Both the bending modulus of the membrane and the elastic area compressibility modulus can be recovered. The bending modulus κ reflects the propensity of the membrane to exhibit curvature. The elastic area compressibility, *K*_*A*_, characterizes the increase of membrane tension δ*T* required to expand a membrane of area *A* by a small increment δ*A*:

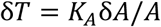

In a former attempt, we had injected septins in the vicinity of the GUVs after inducing micropipette aspiration (Beber *et al*., 2019b) with the initial tongue not in contact with septins. In this configuration, septin binding results in differences in coverage between the aspirated tongue and the outer membrane, leading to a partially heterogeneous distribution. As this may influence the mechanical measurements, we modified the protocol by pre-incubating GUVs with septins prior to aspiration, thereby ensuring a more homogeneous septin coverage over the entire membrane. GUVs were then aspirated with the initial septin density being homogeneous on the surface of each individual GUV.

Without septins, GUVs were aspirated and membrane tongues were induced by micropipettes from the lowest pressure step achievable with our setup (10 Pa). By contrast, in the presence of septins a minimal pressure of about 30 Pa was required to start aspirating a membrane tongue into the pipette (Figure 1.A). Figure 1 displays GUVs with low septin density (A) or three times higher protein density bound to the membrane (B). Upon increasing the aspiration pressure up to 90 Pa, the tongue length remained unchanged until reaching a second critical pressure of about 100 Pa, above which the tongue elongates rapidly. As visualized in Figure 1, the septin signal is absent from the membrane inside the micropipette for both conditions, indicating that septins are excluded from the aspirated membrane tongue. In contrast, the septin fluorescent signal and thus the septin density bound to the membrane outside the pipette increases. These observations suggest that the septin filamentous network is strongly bound to the membrane, and is strongly disrupted and altered to allow for the aspiration of septin-less membrane into the micropipette. For GUVs with higher initial septin density (Figure 1.B), we qualitatively observed that higher pressures were needed both for the initiation of vesicle aspiration and to perform complete vesicle aspiration (see Figure 1.B). Taken together, these observations suggest that septins stiffen membranes. We speculate that at low aspiration pressures, the septin scaffold resist to deformation of the membrane. Because the vesicle volume remains constant, the tongue cannot elongate. Above a critical pressure, the stress exerted becomes sufficient to locally disrupt the septin mesh, so that bare membrane can be aspirated..

**Figure 1.**
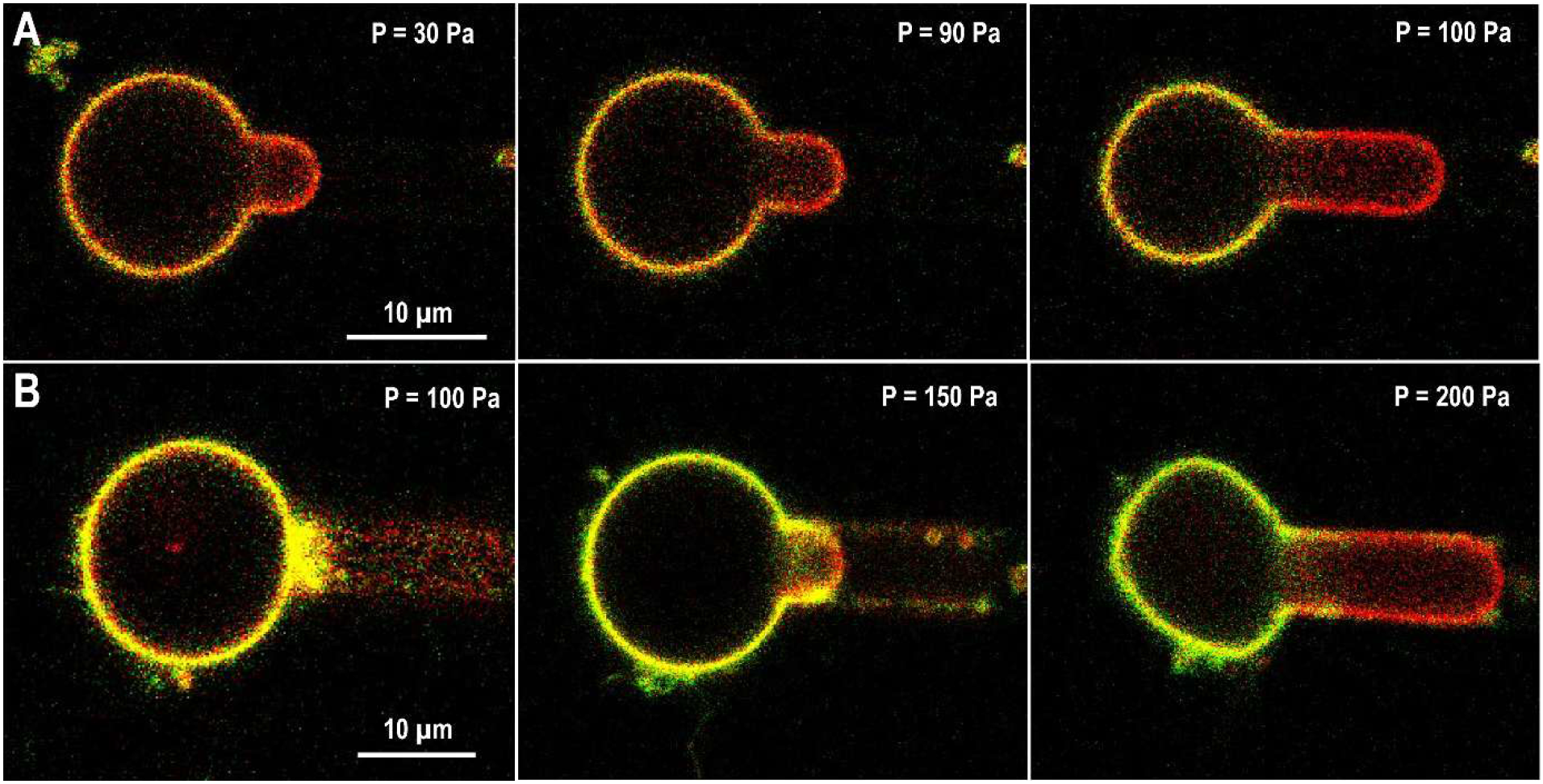
Confocal images obtained during micropipette aspiration of GUVs with a lipid marker (red) in the presence of septins 200 nM (green). In row B the initial septin density on the membrane is 3 times higher than in row A.

These experiments qualitatively describe the role of septins when bound to a vesicle. However, because proteins are excluded from the micropipette, extracting quantitative data from those assays would be misleading. Indeed, bending moduli would usually be estimated from similar assays (Shi and Baumgart, 2015) but require the protein of interest to be homogeneously partitioned on the membrane. To get quantitative insights, we undertook AFM indentation experiments to measure how septins alter membrane deformability.

### AFM Indentation on GUVs : Septins reduce stretching modulus

To perform AFM indentation experiments, spherical colloidal tips were connected to a cantilever to apply a controlled pressure downwards, onto a vesicle. The vesicle deforms in response to the cantilever deflection (Figure 2.A). The force and the vertical position of the tip were extracted from the cantilever deflection and position. The deformation of the vesicle imposed by the tip can be recovered using several models. A Hertz contact approach (Hertz, 1882) was developed to model the contact between two spheres with distinct radii and Young’s moduli. From this model, the Young’s modulus of adherent cells (Krieg *et al*., 2019) or thin materials such as model membranes on a substrate can be assessed (Butt and Franz, 2002b; Redondo-Morata *et al*., 2016). Unfortunately, vesicles are non-homogeneous fluid spheres, and therefore the Hertz model cannot be rigorously applied to determine the mechanical properties of their lipid bilayers. Instead, we employed a model developed in (Schäfer *et al*., 2015), which exclusively considers the tension and area stretching elasticity of a fluid sphere. The model neglects the membrane’s bending modulus, which is appropriate over the length scales involved in our assay. The exact shapes of vesicles are predicted for a given force, allowing the prediction of the indentation depth at this specific force (see Figure 2.A, right). Experimental data were fitted by the model prediction to recover the area stretching modulus (Figures 2.B-C).

**Figure 2.**
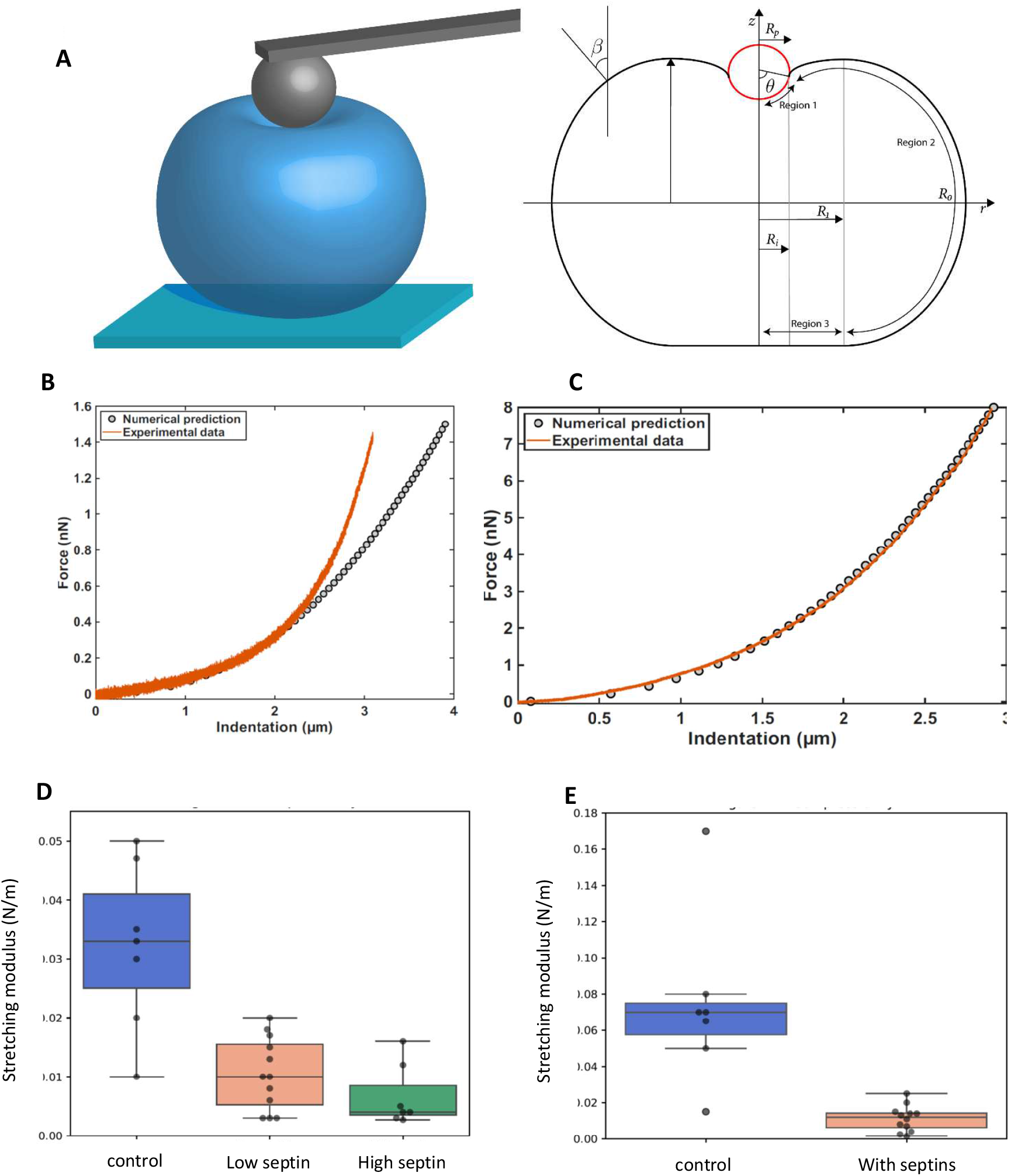
A: (left) Schematic representation of the indentation experiment. (right) Representation of the model from Schäfer (Schäfer *et al*., 2015). The shape of the vesicle is recovered at a known applied force and from this the position of the tip and therefore the indentation. B: Experimental AFM indentation curve (orange) using a tip of radius 1.7 µm and the prediction from the Schäfer model (black circles). For indentations larger than the tip radius, the measurements deviates from the model predictions as shown here. C: Experimental AFM indentation curve (orange) using a tip of radius 5.1 µm and the prediction from the Schäfer model (black circles). D: Distribution of the elastic area stretching for individual vesicles using a tip with a radius of 1.7 µm. E: Distribution of the elastic area stretching for individual vesicles using a tip with a radius of 5.1 µm.

GUVs of 1 to 10 µm in diameter were prepared by gel-assisted swelling (Weinberger *et al*., 2013). Spherical tips of radii 1.7 or 5.1 µm were used to indent the GUVs. Vesicles were immobilized on the surface using different strategies (see methods). Throughout the AFM indentation experiment, vesicles were simultaneously imaged with wide field fluorescence imaging to detect any deformations and lateral displacements (see suppl. Movies 1 and 2). From the resulting observations we could monitor whether the interactions of the GUVs on the glass coverslip were strong enough to prevent any displacement (Supplementary movies 1 and 2). Any experiment where GUV displacement was observed was discarded. Consecutive cycles of indentation were performed on each vesicle, increasing gradually the maximal force applied from 1 to 10 nN. Performing consecutive cycles optimized the position of the tip to the center of the vesicle at low forces and allowed unfolding of potential excess membrane. Besides, it provided force indentation curves with maximal available force before visualizing any rupture of the vesicle.

Spherical AFM probes with a radius of 1.7 µm were used to indent GUVs. The area stretching modulus decreased with increasing septin concentrations, (Figure 2.D), indicating that membranes become easier to stretch upon septin binding. The vesicles had a stretching modulus of 32 ± 6 mN/m (N=7 GUVs) in the control experiment, 11 ± 2 mN/m (N=12 GUVs) in the presence of 150 nM of septins and 7 ± 2 mN/m (N=7 GUVs) in the presence of 300 nM of septins. The analysis could only be performed for indentations smaller than the radius of the tip. Typically, the force corresponding to indentations of 1.7 µm ranged from 1 to 3 nN. For indentations larger than the tip radius, the measurements would deviate from the model predictions as shown in Figure 2.B, mainly due to the enhanced contact between the cantilever and the GUV. To investigate whether GUVs behaved differently under higher applied forces, we used larger beads with a radius of 5.1 µm (Figure 2.C) which allowed to use forces up to 10 nN, corresponding to indentations of less than 5 µm. Again, the presence of septins reduced the stretching modulus of the vesicles from 74 ± 20 mN/m (N=7 GUVs) to 11 ± 2 mN/m (N=12 GUVs) (see Figure 2.E). This drop in stretching moduli reflects the fact that the membrane can be more easily extended in the presence of septins. Taken together, our observations were unexpected, as one might have anticipated that a septin filament network bound to the membrane would reinforce it and increase its stiffness.

### Septins induce reshaping of GUVs prior to indentation

Upon incubation with septins, vesicles appeared deformed prior to indentation, as already visualized in previous studies (Beber *et al*., 2019b; Nakazawa *et al*., 2022). The typical dimensions of the wavelength and the amplitude of deformations were in the micrometric range. The amplitude of deformations was septin dependent. At 150 nM septin concentration, small and regular micrometer deformations were observed. In the presence of 300 nM septins, the amplitude of the deformations was more pronounced, showing sharp bends (see Figure 3.A). Fluorescence images of the GUVs in the presence of septins are displayed in Figure 3.A. The density of septins on the membrane was uniform and appeared homogeneous with no specific enrichment within highly curved or deformed domains.

**Figure 3.**
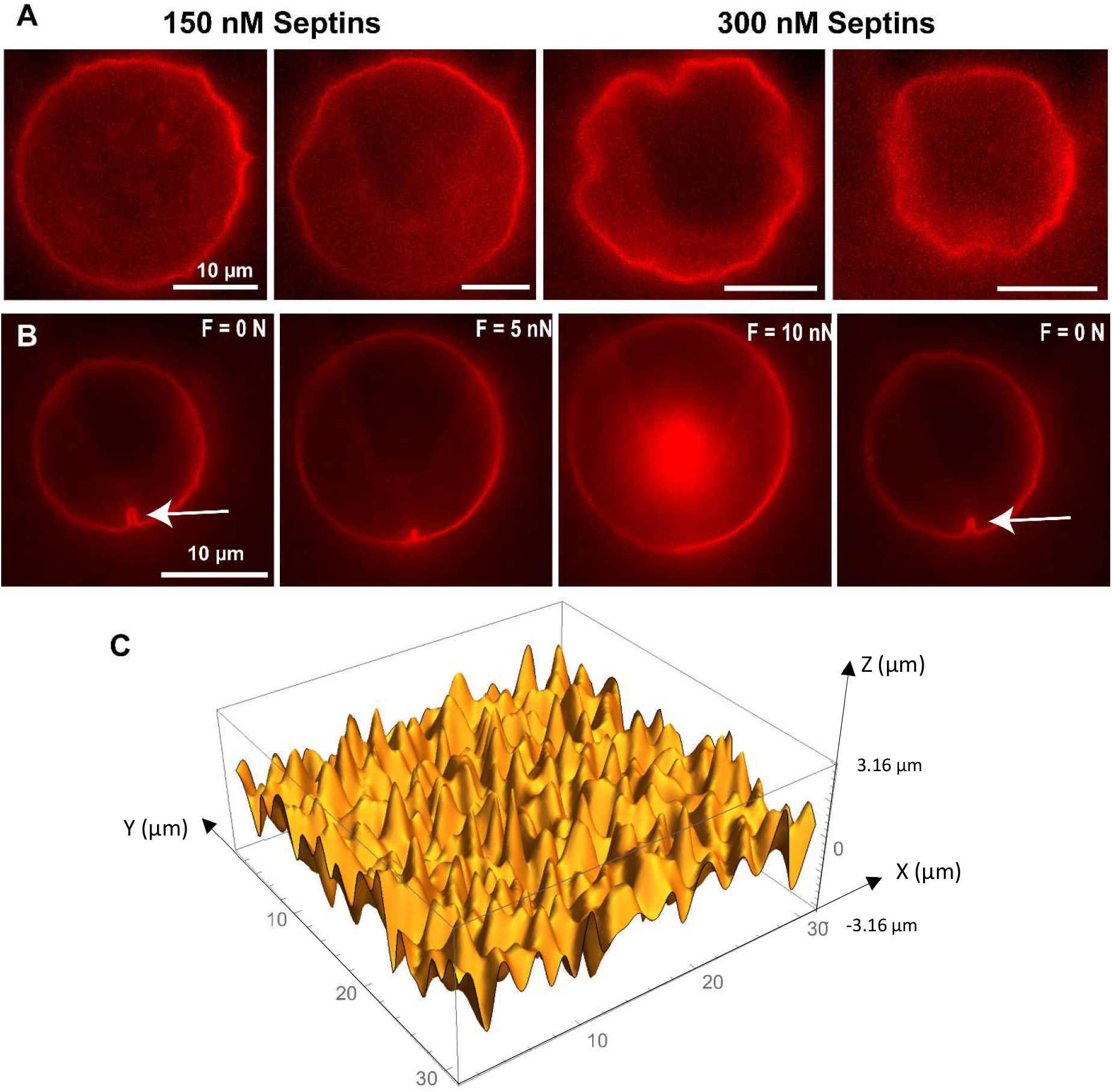
A: Fluorescence images of vesicles with a lipid marker and incubated with different septin concentrations. B: Snapshots from a AFM indentation experiments showing an initial defect (arrow) gradually unfolding and recovered after indentation. C: Realization of a simulated surface following the distribution of curvature used as intrinsic curvature used to model the vesicles, as exposed in the manuscript. Note the uneven horizontal and vertical scales. The former is in units of *a* and the latter in units of *a*^2^*B*^1/2^.

Snapshots from a representative movie (supplementary movies 1 and 2) acquired during a vesicle indentation experiment in the presence of septins are displayed in Figure 3.B. Once a pressure was applied by the AFM spherical probe, the membrane deformations gradually unfolded until the membrane recovered a regular smooth shape similar to the one observed in a control experiment. After the tip was retracted to release the pressure, the deformations were instantly recovered and the shape of the vesicle would perfectly match its initial shape, prior to indentation. This suggests that membrane deformations induced by the septin network are fully reversible over large pressure variations. The deformations and thus membrane folds could thereby provide a membrane reservoir recovered following unfolding. To investigate this hypothesis, a model considering the micrometer membrane deformations was tested to describe our experiments.

### Modeling: membrane deformations to stretching modulus decrease

To understand whether the initial deformations could account for the apparent reduction in stretching modulus, we hypothesized that the presence of the septin network could locally induce an intrinsic membrane curvature. We implemented this feature in the standard Helfrich formalism through a randomly distributed spontaneous curvature. The resulting Hamiltonian can be used to compute the shape of the membrane as a function an externally imposed tension *σ*:

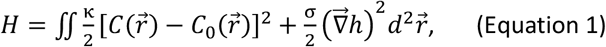

where *κ* is the bending modulus of the thin layer formed by the membrane interacting with septins. We hereafter refer to this composite layer as the “membrane” for simplicity. Equation 1 uses the Monge gauge, as is appropriate for relatively small deformations and in the absence of membrane overhangs. We use these assumptions throughout. We parametrize the membrane shape by the function 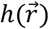 which characterizes its vertical deviation from a perfectly flat shape at position at position 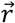. We define its local curvature as 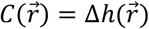, and denote its spontaneous septin-induced curvature at position 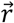 by 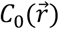. We modeled this spontaneous curvature as a Gaussian colored noise, as illustrated in Figure 3.C. This random field is characterized by two parameters: the typical magnitude *B* of the curvature and the length scale *a* characteristic of the lateral extent of a curvature fluctuation (see the method section for details).

The deformations due to the curvature generate some excess surface that is available upon membrane unfolding. In our description the excess surface per unit membrane surface area reads (see methods):

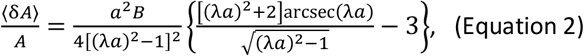

where 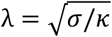 is the inverse of the characteristic length scale associated with the competition between the membrane bending modulus and tension. In all our experimental conditions this length was much larger than the typical distance *a* between curvature peaks, namely λ*a* ≫ 1. In that regime, we can approximate the relative change of area as

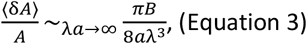

Taking advantage of this observation, we obtain an apparent stretching modulus:

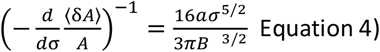

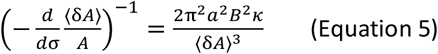

which clearly describes the experimentally observed decrease of the membrane stretching modulus upon an increase of the amplitude *B* of the septin-induced deformations. This modulus diverges as these deformations vanish. In practice, this divergence is prevented by other mechanisms not considered here, *e*.*g*., thermal fluctuations of the membrane, that come to dominate for very small values of *B*. To obtain an independent estimate of *δA*, we considered the geometry used by us in a previous study of the curvature preference of septins (Beber *et al*., 2019b), namely a 2D sinusoidal surface. In such a geometry, the initial excess of membrane is ⟨*δA*⟩ = 0.3. We chose *a* = 1 µm and *B* = 10^13^ m^-2^ to depict membranes with geometrical parameters similar to the ones used in (Beber *et al*., 2019b). Finally, we estimated the bending modulus of the membrane can be estimated from the typical lateral spacing *l* between septin filaments and the persistence length of each individual filament:

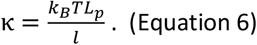

Here *k*_*B*_*T* is the thermal energy and *L*_*p*_ is the persistence length of septin filament. The persistence length of septin filaments has been derived experimentally in various studies. We chose an average value of 10 µm. A typical distance of *l* = 20 nm between filaments was extracted from SEM and TEM images of septin filaments on lipid bilayers (Beber *et al*., 2019b; Nakazawa *et al*., 2022). This yielded *κ* = 2 × 10^−18^ N.m, about 25 times higher than the typical bending modulus of a lipid bilayer. We therefore neglected the contribution of the lipids to κ.

Using these parameter values we obtained an apparent stretching modulus of 10 mN.m^-1^. This value is in close agreement with the experimental value of stretching modulus obtained for GUVs in the presence of septins (Figure 2.B,D,E). From this model, we can conclude that the micrometric deformations induced by septins play a key role in the mechanical behavior of vesicles in the course of indentation.

### AFM indentation on small and non-septin-deformed GUVs

To confirm that the reduction in apparent stretching of the vesicle was exclusively the consequence of initial deformations of the vesicles, we probed the mechanical properties of vesicles smaller than the typical size of the deformations imposed by septins, which thereby appear spherical, initially. If any, deformations affecting those small vesicles would be below the range of deformations observed on larger GUVS and are not visible, with the available resolution. To assess whether vesicles that were not deformed in the micrometer range experienced a reduction in their stretching, we used GUVs with radii smaller than the typical size of the deformation (below 3 µm). AFM tips with beads displaying a radius of 1.7 µm were used. In these conditions, we had to adapt the model proposed in (Schäfer *et al*., 2015) to smaller GUVs. In conditions where the bead is larger than vesicle that is being indented, the Schaffer model becomes numerically unstable and no reasonable solution could be recovered. Therefore, we used the Hertz model. As opposed to the results obtained using larger vesicles, small GUVs display a higher Young’s modulus in the presence of septins (Figure 4.B). Hence, septins rigidify undeformed small GUVs and prior micrometric deformations observed in larger GUVs do play a key role in the vesicle ability to deform. It allows the recovery of a Young’s modulus for an elastic ball. In the presence of 300 nM of septins, small GUVs appeared spherical prior to indentation throughout their imaging by fluorescence imaging (see Figure 4.A). The distribution of apparent Young’s modulus for each single vesicle is displayed in Figure 4.B. The addition of septins caused an important increase in the Young’s modulus of the vesicles from 3.8 ± 0.5 kPa (N=13 GUVs) to 12 ± 2 kPa (N=12 GUVs). The distribution of elastic area stretching modulus as a function of the vesicle radius is displayed in Figure 4.C.

**Figure 4.**
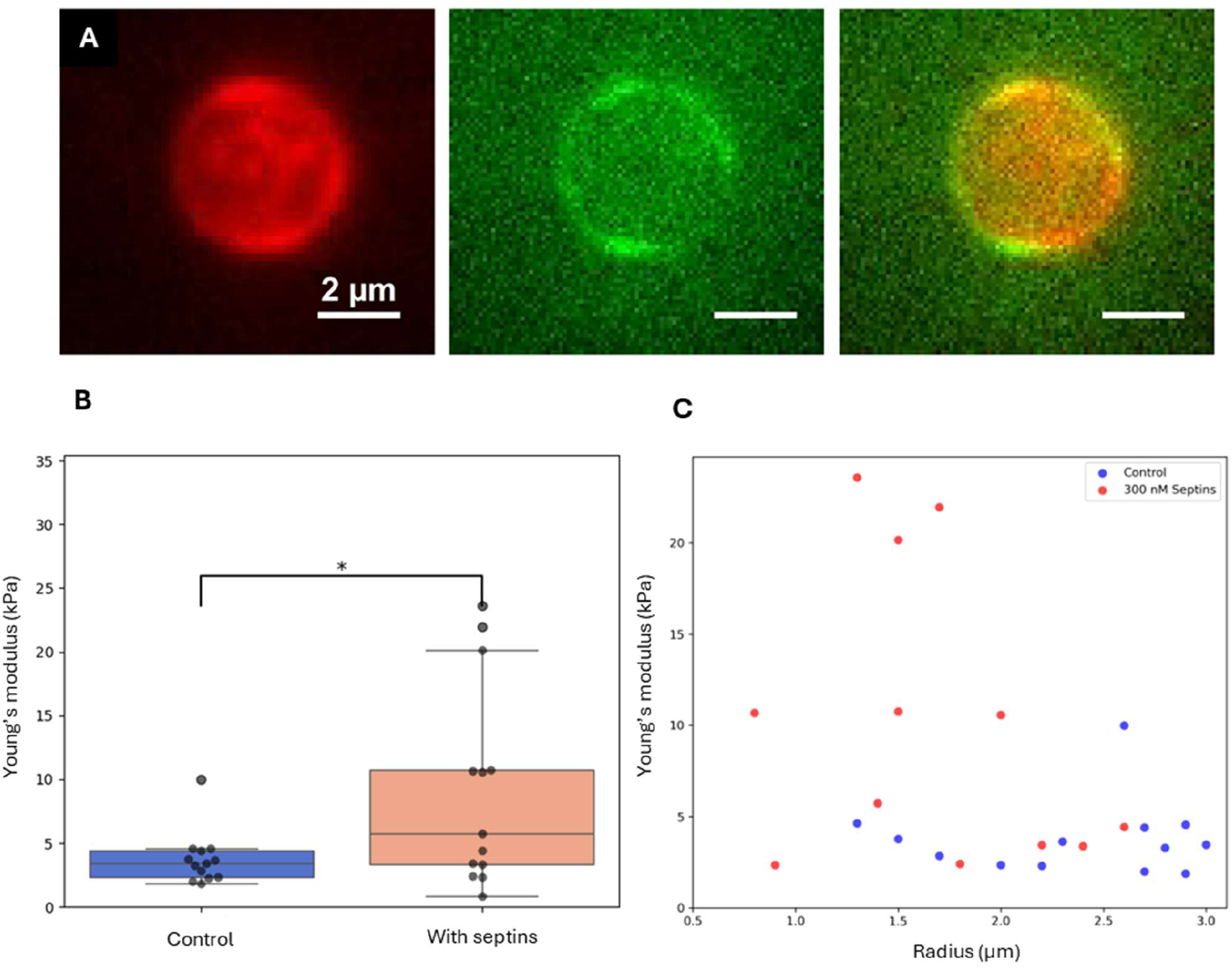
A: Fluorescence image of a GUV marked with a lipid marker (red) prior to indentation in the presence of 300 nM of septins (green). B: Distribution of apparent Young’s moduli modulus of individual GUVs. C: Distribution of apparent Young’s moduli as a function of the vesicle radius.

Taken together, our observations show that, counterintuitively, micrometric membrane deformations induced by septins make GUVs more deformable. However, non-deformed small vesicles when decorated by septins become more rigid.

## Discussion

When measuring the mechanical properties of biomimetic membranes, different assays can be considered. Using GUVs, the micropipette aspiration method developed by Evans (Kwok and Evans, 1981) estimates the fluctuations of stretching moduli by assessing fluctuations in the membrane surface area. This method is valid as long as the membrane remains homogeneous outside and within the micropipette. With septins bound, it is not clear whether septins do indeed enter into the membrane tongue, aspirated in the pipette. When performing those experiments (Beber *et al*., 2019b), the outcome was rather counterintuitive, suggesting that septins barely affect the bending modulus of membranes. AFM spectroscopy probes the global mechanical properties of the septin cytoskeleton bound to a membrane. AFM spectroscopy can be used to assess the mechanical properties of supported lipid bilayers as well (Saavedra V *et al*., 2020). However, as observed previously, septins dramatically disrupt lipid bilayers deposited on mica (Vial *et al*., 2021) and the lipid-substrate interaction can possibly alter the findings. In addition, mechanical properties measured from septins bound to supported lipid bilayers, using AFM, would probe the local organization of septins and may rather reflect the strength of the interaction between septins and the membrane.

Pipette aspiration assays (Figure 1) tend to qualitatively show that a scaffold of septins can stiffen vesicles while AFM indentation experiments suggest that septins make GUVs more deformable (Figure 2). This discrepancy most likely results from our experimental conditions. Indeed, our analysis suggests that septins initially induce micrometric deformations and thus create a membrane reservoir within the resulting membrane folds. Hence, the mechanical constraints applied by an AFM tip unfolds the membrane without being disturbed by the rigidity of the cytoskeletal septin scaffold. When performing the pipette aspiration experiment, the membrane is held with a pipette which imposes a membrane tension, preventing the septins from reshaping the membrane and inducing a substantial membrane reservoir.

Our bottom-up assay might reflect the behavior of membranes *in situ*. Indeed, septins are often associated with membrane remodeling events for which dynamic membrane plasticity is required. For instance, membranes have to be reshaped during macropinocytotic or endocytotic events (Gilden and Krummel, 2010) on different time scales. Septins are known to localize specifically at micrometric curvatures (at the base of cilia, dendrites, at the annulus of spermatozoa or at division furrows) (Cannon *et al*., 2017). In addition, they can display a variety of organizations when bound to membranes. They can self-assemble into parallel sets of filaments that are either parallel or perpendicular to the main curvature axis (Vrabioiu and Mitchison, 2006; Bertin *et al*., 2012). They are also found as orthogonal networks of filaments (Ong *et al*., 2014; Nakazawa *et al*., 2022). Curved membranes favors a specific septin ultrastructural organization, which might, in turn, induce local membrane deformations. Such deformations might thus facilitate membrane reshaping in a cellular context, in synergy with other active factors. Conversely, at shorther lengthscales, alternative septin organizations might favor a global stiffening of membranes required to enhance the stability of a given cellular structure. Hence, septins could be viewed as adaptable proteins that can either reshape or stabilize membranes in a curvature-dependent manner depending on their local ultrastructural organization.

## Material and methods

### Reagents and material

Common reagents (ethanol, acetone, chloroform, sucrose, sodium chloride, Tris) were purchased from VWR reagents and Sigma-Aldrich Co.. L-α-phosphatidylcholine (EggPC, 840051P), cholesterol (700000P), 1,2-dioleoyl-sn-glycero-3-phosphoethanolamine (DOPE, 850725P), 1,2-dioleoyl-sn-glycero-3-phospho-L-serine (DOPS, 840035P), and L-α-phosphatidylinositol-4,5-bisphosphate (PI(4,5)P2, 840046P) were purchased from Avanti polar lipids. Bodipy-TR-ceramide was purchased from Invitrogen (D-7540).

### Septin Purification

Yeast septins were purified from E.Coli extracts following protocols described previously (Bertin *et al*., 2008). Derivatives of pACYCDuet-1 (EMD Biosciences) and pETDuet-1 (EMD Biosciences) carrying respectively cdc3, cdc11 and GFP-cdc10, (His)_6_-cdc12 were co-transformed into E.Coli BL21(DE3) by heat shock. The resulting co-transformants were plated on an agarose gel containing LB medium, chloroamphenicol and ampicilin. Colonies were then picked and were left to grow at 37°C in LB medium containing 34 mg/mL of chloroamphenicol and 50 mg/mL of ampicilin until they reached an optical density of A_600nm_= 1. They were then induced with 500 mM of IPTG for 20h at 16°C. The cells were then harvested by centrifugation and resuspended in Lysis Buffer (300 mM NaCl, 2 mM MgCl_2_, 40 mM GDP, 1 mM EDTA, 5 mM β-mercaptoethanol, 0.5% Tween-20, 12% glycerol, 50 mM Tris, Ph = 8). Cells were lysed using combination of lysozyme treatment (0.5mg.mL^-1^) and sonication in the presence of anti-protease and benzonase. The obtained lysate was centrifuged at 10,000 g for 30 min at 4°C. The septin complexes in the supernatant were captured by Ni^2+^ affinity using Ni-NTA agarose beads (Qiagen). After elution with 0.5 M imidazole, 0.5 M NaCl, 50 mM Tris (pH = 8), the eluate was desalted by passage into a PD-10 column (GE Healthcare) to a buffer containing 150 mM NaCl, 50 mM Tris (pH = 8). Two further steps of purification were performed. First, a size exclusion chromatography was performed using a Superdex 200 column (GE Healthcare) in the same buffer as previously. Then the fractions containing the complexes were pooled and submitted to anion exchange chromatography. To perform this second chromatography, the pooled fractions were first desalted using a PD-10 column into a 50 mM Tris (pH = 8) buffer. They were then injected into a Resource Q anion exchange column (GE Healthcare) and submitted to a linear salt gradient, starting from 0 to 0.5 M of NaCl. Typically, septin complexes eluted around 300 mM of NaCl. This concentration of salt prevents further polymerization of the complexes. The fractions containing septins were aliquoted, flash frozen and kept at -80°C for later use. The concentration of the samples was measured using a NanoDrop™. The extinction coefficient for the GFP-labelled octamer is ε = 291.000 M^-1^.cm^-1^. The typical yield was 1 mg per liter of culture. The quality of the proteins was analyzed by running SDS-page gel.

### GUV preparation

The “Growth Buffer” containing 50 mM Sucrose, 50 mM NaCl and 10 mM Tris (pH = 7.8) and “Observation Buffer” containing 75 mM NaCl and 10 mM Tris (pH = 7.8) were prepared. Their osmolarity was measured using a freezing point osmometer (Löser) and adjusted by adding small amounts of NaCl until their difference was less than 5%.

GUVs were prepared using the gel-assisted swelling method. A solution of 5% w/w PVA, 150 mM Sucrose and 10 mM Tris (pH = 7.8) was prepared. To ensure dissolution of the PVA the solution was heated to 95°C for 30 min or until full dissolution. 20 µL of the solution was homogeneously spread on a 22*22 mm coverslip previously rinsed with water and ethanol and dried using nitrogen flow. The coverslips were then placed in an oven at 60°C for 30 min. and later kept at room temperature and could be used for up to two weeks.

Lipids were mixed following compositions used in previous studies. A solution of lipids at a concentration of 2 mg.mL^-1^ in chloroform was prepared using a molar ratio of 56.8% EggPC, 15% cholesterol, 10% DOPE, 10% DOPS, 8% brain-PIP_2_ and 0.2% Bodipy-TR Ceramide. The solution was kept in amber vials at -20°C for up to 2 months. 10 µL of the solution was spread on each coverslip covered with PVA gel and the coverslips were then placed in vacuum for 30 min to allow full evaporation of the chloroform. 1mL of “Growth Buffer” was then added on the coverslip and coverslips were left for 1 hour to allow growth. The GUVs were then collected by first tapping on the bottom of the coverslip to detach GUVs from the gel and then aspirated using a cut pipette tip with an opening of about 2mm.

### Correlative wide-field fluorescence and AFM indentation

AFM experiments were performed on a combination of a Nanowizard 4 microscope (JPK, Bruker) and a in-house developed inverted widefield fluorescence and TIRF microscope equipped with an oil immersion objective (Plan-Achromat 100x, Zeiss) (Dahmane *et al*., 2019; Vial *et al*., 2023). A 1.5x telescope was added to obtain a final 150x magnification and a corresponding pixel size of 107 nm. Two laser lines were used (488 nm and 561 nm, Obis, Coherent) combined into a single beam. The intensity was tuned with an acousto-optic tunable filter (AOTFnc-400.650-TN, AA opto-electronics) and fluorescence images were acquired using a 525/50 nm (Chroma) and 600/50 (Chroma) emission filters for GFP and Bodipy-TR detection, respectively. An ET800sp short pass filter (Chroma) was used in the emission optical path to filter out the light source of the AFM optical beam deflection system, centered at 980 nm. The camera used for imaging fluorescence was an EMCCD iXion Ultra897 (Andor Technologies). The lasers, filters and camera were controlled with a custom made LABVIEW script. The acquisition time for each frame was set at 100 ms. One single image was collected without averaging. To prevent lateral displacement, GUVs were bound to the glass substrate. Circular glass coverslips (2.5 cm, 165 μm thick, purchased from Marienfeld) were used. Coverslips were cleaned with a 15 min cycle of sonication with ultrasounds in 1 M KOH, rinsed 20 times with deionized water and finally with a second cycle of sonication in deionized water. Multiple methods for GUVs binding were used. Strong interactions could result in bursting the GUVs on the glass substrate, leaving only lipid bilayers. Since polylysine left only the smallest vesicle intact, we used a small amount of MgCl_2_ (20 µL) at a concentration of 10 mM that was then diluted and washed away before addition of vesicles which lead to strong binding but very little bursting. A higher concentration leads to bursting of all GUVs within 2 to 3 minutes. GUVs were incubated for 20 min with septins and later injected in the observation chamber. After another 20 min of sedimentation, the chamber was rinsed with the “Observation Buffer” to remove the excess of septins.

AFM indentation experiments were carried out using CP-PNPL-SiO (sQube) triangular cantilevers with nominal spring constant of 0.08 or 0.32 N/m and equipped with spherical colloidal probes with diameters of either 1.7 µm or 5.1 µm. AFM tips were cleaned with air plasma treatment and kept in desiccated atmosphere. The inverse optical lever sensitivity was calibrated with the acquisition of a force versus distance curve on the glass coverslip in absence of GUVs, whereas the cantilever stiffness was calibrated using thermal (Proksch *et al*., 2004).The fluorescence imaging allowed fast scanning of the sample to find GUVs and proper alignment of the AFM tip with the vesicle of interest to indent it at its center. Simultaneous AFM and fluorescence microscopy acquisition required precise control and minimization of optomechanical effects, which necessitated the use of excitation lasers in the microwatt range (Fernandes *et al*., 2020). The vesicles were selected for their unilamellarity and their dimension. When indenting, the force displacement curves were recorded during both the tip approach and the retract to observe if some of the energy was lost by viscosity, and if the tip detached properly from the GUVs. This issue was more important when septins were added as they adhere to the tip. Experiments were performed fixing the maximum force applied by the lever (up to 15 nN): such force was optimized during data acquisition in order to prevent GUVs indentation longer than the spherical probe size. Therefore, different cycles of increasing higher maximum force were performed on each GUV. The speed used for the vertical displacement of the tip was 1µm.s^-1^to ensure quasi-static conditions, whereas the indentation cycle length was fixed to 10 µm to ensure GUV-probe detachment during retract.

The analysis of AFM curves was performed using the JPK software and homemade Matlab routines. The X axis values reported in the indention curves (Figure 2C,D), labelled as “indentation”, are obtained subtracting the position of cantilever base with the cantilever deflection, and therefore represent the contact part of the probe-GUV distance. We used both the model described in Schaffer et al. and the Hertz contact model (Hertz, 1882; Schäfer *et al*., 2015), depending on the vesicle size. The initial radii of the vesicles were recovered from the fluorescence images.

The model proposed by Schäfer et al relies on the Laplace equation to recover the shape of a vesicle on which a force is applied by a tip. From the Laplace equation, three equations can be derived. The first one from the conservation of the volume and the other two from the force equilibrium between the force applied by the tip and the force applied by the membrane on both the tip and on the substrate. Numerically solving this system of equation allows recovery of a predicted indentation depth for a given force depending on the stretching and the initial tension. The experimental data was fitted to obtain the stretching of the vesicle. Because the model relies on solving a system of three non-linear equations it is very sensitive to the initial conditions and can be very unstable depending on the different parameters.

In the Hertz contact model, Because the geometry of the vesicle differs from the geometry considered in the model, the equation relating the force to the indentation does not scale as predicted. Therefore, we considered only the first 200 nm of indentation to obtain an estimation of an apparent Young’s modulus.

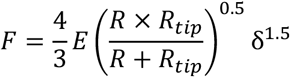

In this model, we consider that the vesicles are elastic half balls instead of a sphere. There are therefore characterized mechanically by a single parameter, their Young’s modulus. Where *F* is the force applied, *E* is the Young’s modulus, *R* is the radius of the vesicle, *R*_*tip*_ is the radius of the AFM tip and *δ* is the indentation depth.

### Micropipette aspiration

A 5 mg.mL^-1^ solution of β-casein was prepared in the “Observation Buffer”. It was used to passivate all glass surfaces to prevent bursting of GUVs on the surfaces due to strong interactions as well as to limit the interaction between septins and glass.

The chamber for observation consisted of two glass coverslips glued with vacuum grease on a spacer with a thickness of 1 mm while the sides were left open. The bottom coverslip was 30 × 11 mm, the top one 35 × 9 mm. The solution of β-casein was injected in the chamber and left for 30 min to fully passivate the glass coverslips.

GUVs were manipulated using glass micropipettes. Micropipettes were made from glass capillaries pulled using a pipette puller P-2000 (Sutter Instruments). They were then forged using a MF-830 microforge (Narishige) to obtain the desired opening size (about one fourth of the diameter of the vesicles corresponding to 5 to 10 µm). They were then filled with the β-casein solution with home made syringes using a thin glass capillary as a needle. The micropipettes were then connected to a water tank which can be moved vertically to control the pressure inside micropipettes. The micropipettes were then fixed on a micromanipulator (Narishige) and inserted inside the observation chamber.

The β-casein solution was removed after incubation and the GUV solution could be added after dilution in the “Observation Buffer”. The ratio for dilution depended on the yield of growth. Typically a ratio of GUV solution to ‘Observation Buffer’ of 1:10 was used. After the sample injection in the observation chamber, the vesicles were left to sediment for 10 to 20 min before their observation. When septins were added, they were first diluted to reach the same osmolarity as in the “Observation Buffer” and mixed with the GUVs just prior to the injection in the observation chamber.

To set the pressure inside micropipette to the value in the chamber, the height of the water tank was adjusted to have no flux of water at the tip of the micropipette. Vertical displacement of the water tank was measured and related to a difference of pressure from inside the micropipette to the observation chamber.

The confocal microscope consisted of an Nikon TE-Eclipse inverted microscope with three laser lines (λ = 488 nm; λ = 561 nm and λ = 647 nm) combined with a Nikon-C1 confocal head.

### Analytical calculations of the membrane area enclosed in the ruffles

The Helfrich Hamiltonian describes the energy of a membrane as a function of its state:

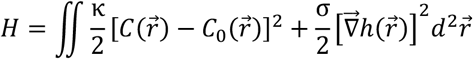

Where *κ* is the bending modulus of the membrane interacting with septins, *C* = *Δh* and *C*_0_ are respectively the local curvature and the intrinsic curvature, *σ* is the tension of the membrane and *h* its height in the Monge gauge. At mechanical equilibrium, the Hamiltonian *H* is minimized with respect to the function 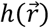, implying that the latter satisfies the following Euler-Lagrange equation:

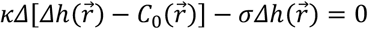

Its Fourier transform thus reads

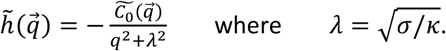

The resting geometry of the membrane is thus determined by the spontaneous curvature field 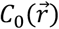. To reflect the apparent random arrangement of septin filaments in our experiments, we modeled it as a colored Gaussian noise. This stochastic process indeed combines a simple analytical structure with the ability to introduce a length scale *a* over which the structure of the septin network is correlated with itself. The distribution of the curvature is fully characterized by its two first cumulants:

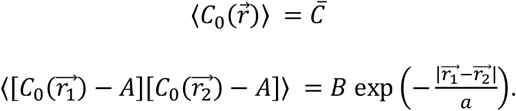

We aim to compute the area contained in the septin-induced deformations, namely:

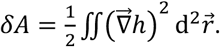

Using the Fourier transform it becomes:

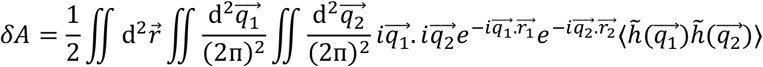

To compute 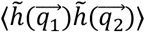 we define the quantity 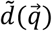 as:

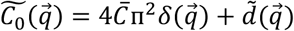

which implies

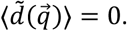

Then:

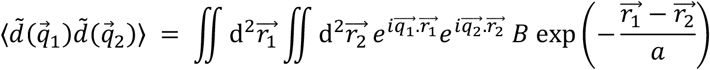

By changing the variable: 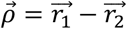 the equation becomes:

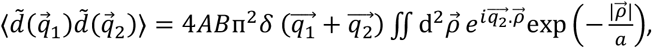

Where *A* is the total projected area of membrane. We choose a system of coordinates whose x axis is aligned with 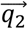 through 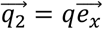 and define the angle *ϕ* of polar coordinates as:

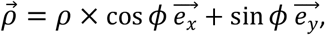

which yields

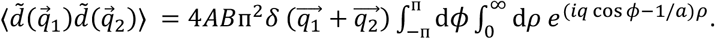

We integrate over *ρ* and obtain:

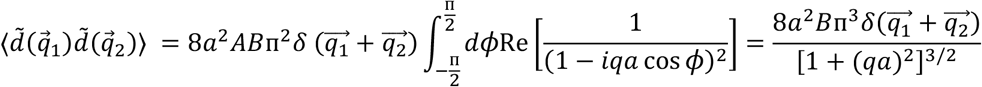

and thus:

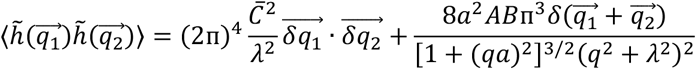

which we use to compute

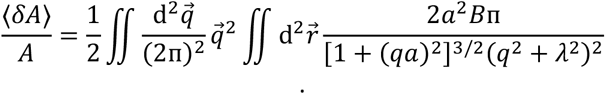

The average of the relative area contained in the septin-induced membrane deformations thus reads:

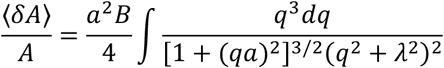

Changing the integration variable to *Q* = (*qa*)^2^, the equations becomes

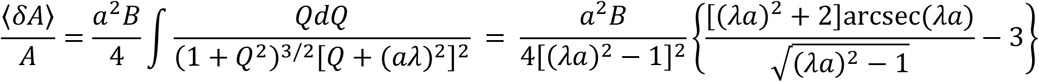

as specified in Equation (2) of the main text. Note that this stored area is independent of the mean spontaneous curvature 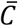: only the curvature fluctuations (quantified by *B*) are relevant.

## Acknowledgements

This work benefited from the support of the ANR (Agence Nationale de la Recherche) for funding the project “SEPTIME”, ANR-13-JSV8-0002-01 and the project “SEPTSCORT”, ANR-20-CE11-0014-01. B. Chauvin is funded by the Ecole Doctorale “ED564: Physique en Ile de France” and Fondation pour la Recherche Médicale. We thank the Labex Cell(n)Scale (ANR-11-LABX0038) and to Paris Sciences et Lettres (ANR-10-IDEX-0001-02). The CBS is a member of the France-BioImaging (FBI) and the French Infrastructure for Integrated Structural Biology (FRISBI), two national infrastructures supported by the French National Research Agency (ANR-24-INBS-0005 FBI (BIOGEN) and ANR-10-INBS-05, respectively). This work was also supported by the GIS IBISA. M.L. was supported by “Investissements d’Avenir” LabEx PALM (Grant No. ANR-10-LABX-0039-PALM), ANR Grants No. ANR-21-CE11-0004-02, No. ANR-22-ERCC-0004-01, and No. ANR-22-CE30-0024-01, as well as ERC Starting Grant No. 677532 and the Impulscience® program from Fondation Bettencourt-Schueller. M.L.’s group belongs to the CNRS consortium AQV.

